# The maternal vGluT2 and embryonic mGluR3 signaling relay system controls offspring wing dimorphism in pea aphid

**DOI:** 10.1101/2024.05.22.595290

**Authors:** Yiyang Yuan, Yanyan Wang, Wanwan Ye, Liqiang Xie, Erliang Yuan, Huijuan Guo, Shifan Wang, Fang Dong, Keyan Zhu-Salzman, Feng Ge, Yucheng Sun

**Author notes:** Author for correspondence: Feng Ge; Yucheng Sun.

## Abstract

Transgenerational phenotypic plasticity (TPP) refers to the phenomenon that environmental conditions experienced by one generation can influence the phenotype of subsequent generations to adapt to the environment without modification of their DNA sequences. Aphid wing dimorphism is a textbook example of TPP by which a maternal aphid perceives the environmental cues to decide the wing morph of her offspring. However, the signaling mechanism from mother to daughter remains unclear. In this study, we showed that the population density and physical contact caused high proportion of winged offspring in the pea aphid *Acyrthosiphon pisum*. Its *vesicular glutamate transporter 2* (*ApvGluT2*) and *metabotropic glutamate receptor 3* (*ApmGluR3*) were identified by tissue-specific RNA-seq as differentially expressed genes in the head and embryo respectively between solitary and more densely housed maternal aphids. Elevated expression of brain *ApvGluT2* and embryonic *ApmGluR3* led to increases in the winged proportion. Knockdown of either gene inhibited phosphorylation of ApFoxO in embryos. Furthermore, EMSA showed that dephosphorylated ApFoxO directly bound to the promotor of *hedgehog* (*ApHh*), a morphogen gene for wing development, to repress its transcription in stage 20 embryos, causing a lower winged proportion. Our results demonstrated that brain *vGluT2* and embryonic *mGluR3* coordinately relayed the maternal physical contact signals and control wing development in offspring, showcasing a novel regulatory mechanism underlying physical contact-dependent, transgenerational wing dimorphism in aphids.

## Introduction

The transgenerational phenotypic plasticity (TPP) is a fascinating capacity of individual genotypes, when experiencing different environmental conditions, to produce different phenotypes in subsequent generations (*Bell and Hellmann, 2019*). Since the impact of parental experience is expected to persist across generations, the phenotypic adjustment warranted by parents could be beneficial to offspring (*Guillaume et al., 2016*). This process involves detection of environmental cues by a parent, signal delivery, and signal perception by offspring and subsequent alternative phenotypes (*Bell and Hellmann, 2019*). Parent-to-offspring signaling can be accomplished via mechanisms that are rapid and non-genetic, such as through neurotransmitters, hormones, epigenetic modifications, and microbiota (*Donelan et al., 2020*).

Insect wing polyphenism is an intrinsic characteristic; a single species can produce flightless (wingless or short-wing) and flight-capable (winged or long-wing) morphs depending on the environment (*Brisson, 2010*; *Zera, 1997*; *Zhang et al., 2019*). Numerous originally winged insect species have become flightless because secondary flight loss favors their adaptation to environmentally stable habitats (*Roff 1990*; *Roff, 1994*; *Wagner and Liebherr, 1992*). Aphid wing dimorphism is an iconic example of TPP in that maternal aphids perceive the environmental cues, such as population density, nutrients, natural enemy, or even physical contact, and signal the offspring during the embryonic period, by which alate (winged) or apterous (wingless) daughters are destined before birth (*Braendle et al., 2005*; *Ishikawa and Miura, 2013*). Of the environmental cues, high population density is widely recognized as a biotic stress reflecting the intraspecific competition within habitat, which could reliably drive the transition from wingless to winged aphids across generations (*Deem et al., 2024*; *Sng and Ackerman, 2020*). Since wingless aphids typically have higher fecundity than the winged morph, switching to the dispersal winged morph from the reproductive morph alleviates resource competition in offspring (*Deem et al., 2024*). In some strains of pea aphids, even two maternal aphids housed in a petri dish can efficiently induce a high proportion (>90%) of winged offspring than singly housed aphids (*Sutherland, 1969*; *Yuan et al., 2023a*; *Yuan et al., 2023b*). Physical contact seems to be a trigger for adults to produce winged offspring (*Sutherland, 1969*). Similarly, the aphid alarm pheromone can also induce groups of aphids, rather than single individuals, to produce a higher proportion of winged offspring, suggesting that the singly housed aphids that lack physical contact cannot initiate wing morph change in offspring despite perception of the alarm pheromone (*Kunert et al., 2005*).

Recent studies have identified a number of genes and gene networks involved in the regulation of insect wing polyphenism. In planthoppers, the zinc finger homeodomain transcription factor (Zfh1) and insulin/insulin-like growth factor signaling (IIS) pathways coordinately modulate the fate of wing development in a Forkhead transcription factor subgroup O (FoxO)-dependent manner (*Xu et al., 2015*; *Zhang et al., 2022*). Dephosphorylated FoxO suppresses wing development by inhibiting the transcription of cellular proliferation genes and wing-patterning genes, or through repressing the transcriptional activity of Rotund, a zinc finger transcription factor of wing developmental genes (*Chen et al., 2023*; *Lin et al., 2020*; *Xu et al., 2021*; *Yuan et al., 2023a*; *Zhang et al., 2021*). However, these mechanisms are within one generation prior to the critical switching time point at the nymphal stage. In contrast, the wing morph of some aphids is determined transgenerationally. Ecdysteroid, miR-9b, dopamine, and IIS signaling in maternal pea aphids are altered by experiencing high population density, which increases the proportion of winged offspring (*Liu and Brisson, 2023*; *Shang et al., 2020*; *Vellichirammal et al., 2017*; *Yuan et al., 2023a*). In offspring, the TOR (the target of rapamycin) and Wnt signaling pathways respectively control the occurrence of autophagic and apoptotic degradation of wing disc in 1^st^ instar nymph of pea aphids, resulting in the formation of wingless morph (*Yuan et al., 2023b*; *Zhou et al., 2023*). However, the molecules in the brain and embryo that are specifically stimulated by maternal density and determining offspring wing morph before birth are largely unknown.

Glutamate is a primary excitatory neurotransmitter. It not only functions in the signal transduction cascade mediating the parent-offspring communication in the locust, but is involved in the regulation of its interconvertible phase change, from the solitarious to gregarious phase in response to high population density (*Yang et al., 2023*). Maintaining an optimal glutamate concentration at the right time and in the right tissue is important for proper nerve cell signaling, and embryo and tissue development. Dysfunctional glutamatergic synaptic transmission has been linked to transgenerational effects on anxiety, fear, pain sensitization, and embryonic development in animals (*Jones et al., 2022*; *Liu et al., 2017*; *Tao et al., 2020*; *Van Winkle, 2022*). Glutamate signaling involves a sophisticated network of metabolic pathways and transport mechanisms that facilitate glutamate synthesis, release, and clearance, ensuring its proper function as the primary excitatory neurotransmitter in the central nervous system (*Featherstone, 2010*). This process begins with glutamate production in the presynaptic neurons via the enzyme glutaminase, followed by glutamate sequestration into synaptic vesicles by vesicular glutamate transporters (vGluTs). Upon synaptic activation, glutamate is exocytosed into the synaptic cleft, where it interacts with ionotropic or metabotropic glutamate receptors (iGluRs and mGluRs) on the postsynaptic neuron, triggering a variety of signaling cascades (*Willard and Koochekpour, 2013*).

The glutamate signaling pathway could modify the phosphorylation level of FoxO. Phosphorylated FoxO no longer binds to promoters and thus loses the suppression of the network of wing development genes (*Howlett et al., 2008*; *Xu et al., 2015*; *Zhang et al., 2018*). It therefore is plausible that the maternal glutamate pathway controls the phosphorylation level the embryonic FoxO, and thereby regulating the developmental trajectory of wing disc in offspring. In this study, maternal *vesicular glutamate transporter 2* (*ApvGluT2*) and *metabotropic glutamate receptor 3* (*ApmGluR3*) of pea aphids were identified by the tissue-specific RNA-seq as differentially expressed genes; both were induced by physical contact or by high population density. Given the importance of the glutamate signaling pathway in perceiving the population density and transducing the signal to offspring, the effects of brain *ApvGluT2* and embryonic *ApmGluR3* were determined on (i) increases in the proportion of winged offspring, and (ii) FoxO phosphorylation level and downstream wing developmental genes at stage 20 embryo stage, the crucial time point for determination of the wing morph.

## Results

### Physical contact and high maternal density increased the proportion of winged offspring

To determine the transgenerational effects of maternal density and physical contact on the wing morph in pea aphid offspring, maternal aphids in groups of 1, 2, and 8 were placed in a petri dish respectively, mimicking the solitary, physical contact, and maternal density effects (Figure 1A). Maternal aphids treated with physical contact and high density for 2- and 4-hr had higher proportions of winged offspring than solitary aphids (Figure 1B). Notably, 2- and 4-hrcontacts between two adults resulted in 54.1% and 96.5% winged offspring, similar to the effects of high-density treatment for the same time periods, 86.5% and 94.6% winged offspring, respectively (Figure 1B). Thus, physical contact could be utilized to obtain effective induction of winged offspring and even explain the effect achieved by high population density.

**Figure 1.**
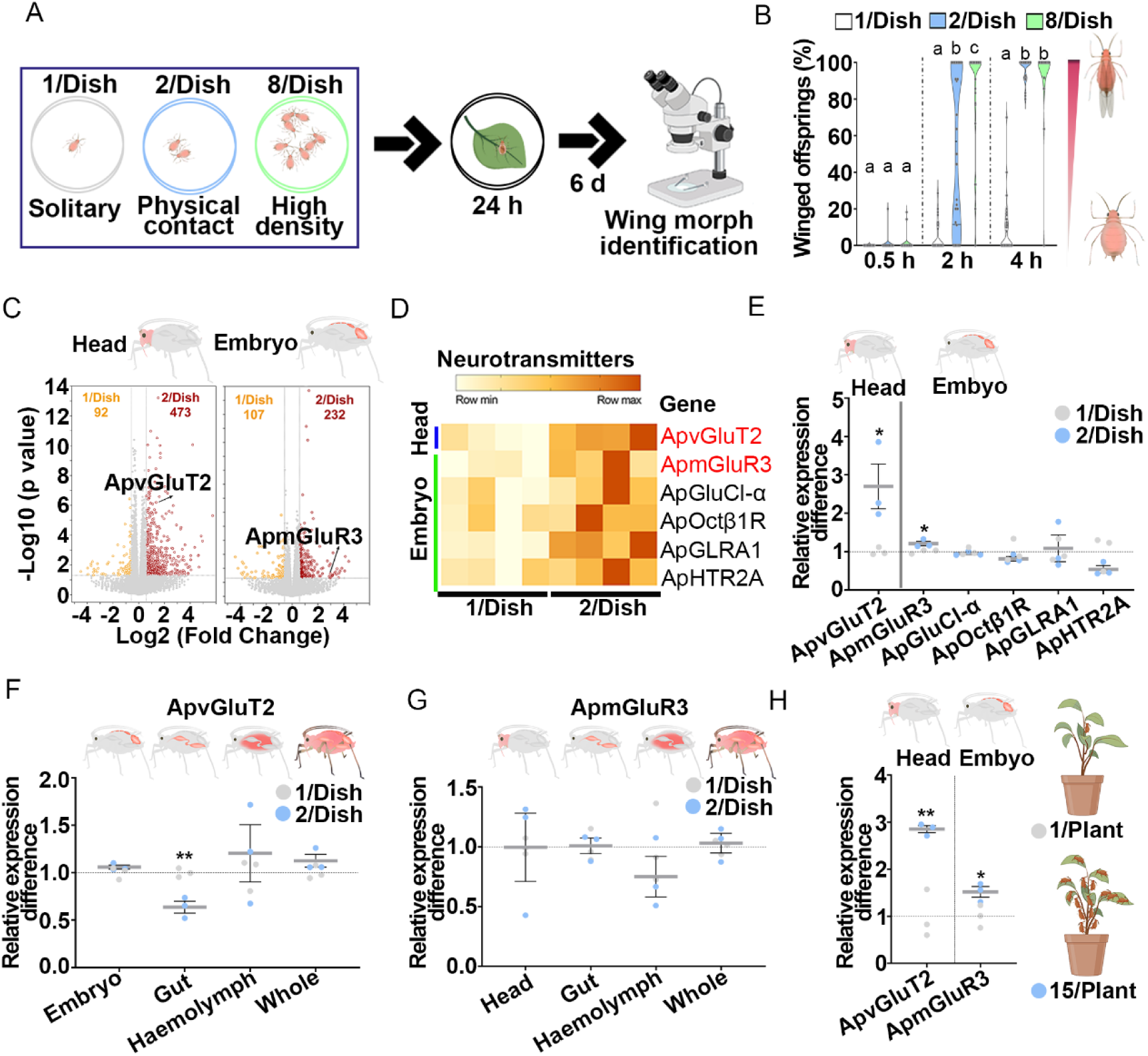
Maternal aphids produced high proportions of winged offspring after experiencing physical contact and high density, with increased expression of head *ApvGluT2* and embryo *ApmGluR3*. (A) The schematic procedure of transgenerational induction of winged offspring. (B) Effects of maternal density and duration on the proportion of winged offspring (n=32). Two-adult contact for 4 hr was sufficient to induce over 90% of winged offspring. Kruskal-Wallis test was performed, and Dunn’s test was used to differentiate between the means; different letters indicate significant differences (P < 0.05). (C) The volcano plots showing differentially expressed genes (DEGs) in the head and embryo induced by two-adult contacting for 4-hr (n=4). DEGs were selected if the absolute value of Log2 (Fold change) > 0.58 and P < 0.05. (D) The heat map showing the expression patterns of six DEGs functioning in neurotransmitter signaling in RNAseq analyses (n=4). (E) qPCR confirmation of increased expression of *ApvGluT2* and *ApmGluR3* in head and embryo, respectively caused by maternal physical contact (n=3). (F) The tissue-specific expression of *ApvGluT2* and (G) *ApmGluR3* in two-adult contacting for 4-hr treatment vs. solitary treatment (n=3). (H) Transcripts of head *ApvGluT2* and embryo *ApmGluR3* were higher in group aphids of 15 adults than those of singly kept aphids on plants, as determined by qPCR (n=3). Student *t* test: *P < 0.05, **P < 0.01.

Since data collected from the two-adult contacting for 2-hr treatment exhibited a full range in percentage of winged offspring from 0% to 100%, the locomotor activity of maternal aphids was traced to determine the possible correlation with the proportion of winged offspring. Three locomotor parameters, including movement distance, velocity, and contact frequency of maternal aphids were monitored by the EthoVision system (Noldus) (Figure S1 A). All three maternal locomotor parameters exhibited strong positive correlations with the winged offspring proportion (*P* value < 0.0001) (Figure S1 B-D). Furthermore, since movement distance and velocity of maternal aphids were not significantly different between solitary and two-aphid contact for 4-hr treatments, physical contact between maternal aphids could be the key driving factor for higher proportion of winged offspring (Figure S1 E-F).

### Physical contact and high density increased the expression of brain *ApvGluT2* and embryonic *ApmGluR3* in maternal aphids

RNA-seq was used to determine tissue-specific gene profiles of solitary maternal aphids and those subjected to 4-hr physical contact treatment. A total of 563 differential expressed genes (DEGs) were identified in maternal heads, while 339 genes were differentially expressed in embryos with the criteria of the absolute value of Log_2_ (fold change) > 0.58 and *P* value < 0.05 (Figure 1C). Of which, only 6 DEGs functioning in neurotransmitter signaling were identified, including *ApvGluT2* (LOC100168565), *ApmGluR3* (LOC107882629), *Glutamate-gated chloride channel α* (*ApGluCl-α*, LOC100162577), *Octopamine receptor β-1R* (*ApOctβ1R*, LOC100166522), *Glycine receptor subunit α1* (*ApGLRA1*, LOC100169196), and *Serotonin receptor 2A* (*ApHTR2A*, LOC100575043) (Figure 1D). qPCR results further confirmed that only head *ApvGluT2* and embryonic *ApmGluR3* had higher transcripts in maternal aphids experiencing physical contact than in solitary aphids (Figure 1E). Genes associated with glutamate signaling components were annotated, and their expression levels in maternal brain and embryo tissues were listed (Figure S2). Furthermore, differential expression of *ApvGluT2* and *ApmGluR3* were not observed in other tissues or in whole body except for gut *ApvGluT2* (Figure 1F-G). To validate the involvement of *ApvGluT2* and *ApmGluR3* in perceiving high population density, 15 adults vs. 1 adult were clipped into a plant seedling for 24-hr, and heads and embryos were then dissected and sampled. RT-qPCR showed that 15-adult treatment increased *ApvGluT2* transcripts by 64.9% in the head and *ApmGluR3* transcripts by 34.2% in the embryo relative to solitary aphids (Figure 1H). These results suggested that maternal brain vGluT2 and embryonic mGluR3 were involved in the physical contact-induced production of winged offspring.

### Maternal *Ap*v*GluT2* expressed in antennal lobes of aphid brain favored winged offspring in a locomotor activity-independent manner

Previous research showed that neural vGluT2 could modulate animal locomotor activity (*Roccaro-Waldmeyer et al., 2018*). It was therefore speculated that physical contact-induced *ApvGluT2* in aphid brain may increase the proportion of winged offspring by enhancing the locomotor activity of maternal aphids or by cascading the glutamate pathway to regulate the wing development across generations. To experimentally test these hypotheses, pharmacological assays were performed on maternal aphids (Figure 2A), followed by monitoring of the locomotor activities and proportion of winged offspring of aphids subjected to 2-adults contact for 2-hr (Figure 2A). Application of glutamate agonist glutamic acid increased winged offspring from 47.6% to 68.1% (Figure 2B). Conversely, vGluT2 inhibitor Chicago Sky Blue (CSB) reduced the proportion of winged offspring from 53.3% to 31.2% (Figure 2F). However, either had any effects on locomotor activities (Figure 2C-E, 2G-I). Fluorescence *in situ* hybridization (FISH) revealed that *ApvGluT2* was mainly expressed in the neural antennal lobes of the aphid brain, and the fluorescent signal intensified as the maternal population density or contact duration increased (Figure 2J-K). Amputating the antennae of maternal aphids significantly reduced the proportion of winged offspring from 95.2% to 77.0% (Figure 2L). Together, aphids activated brain *ApvGluT2* in response to maternal density and physical contact, yet the resulting higher proportion of winged offspring had little to do with the locomotor activity.

**Figure 2.**
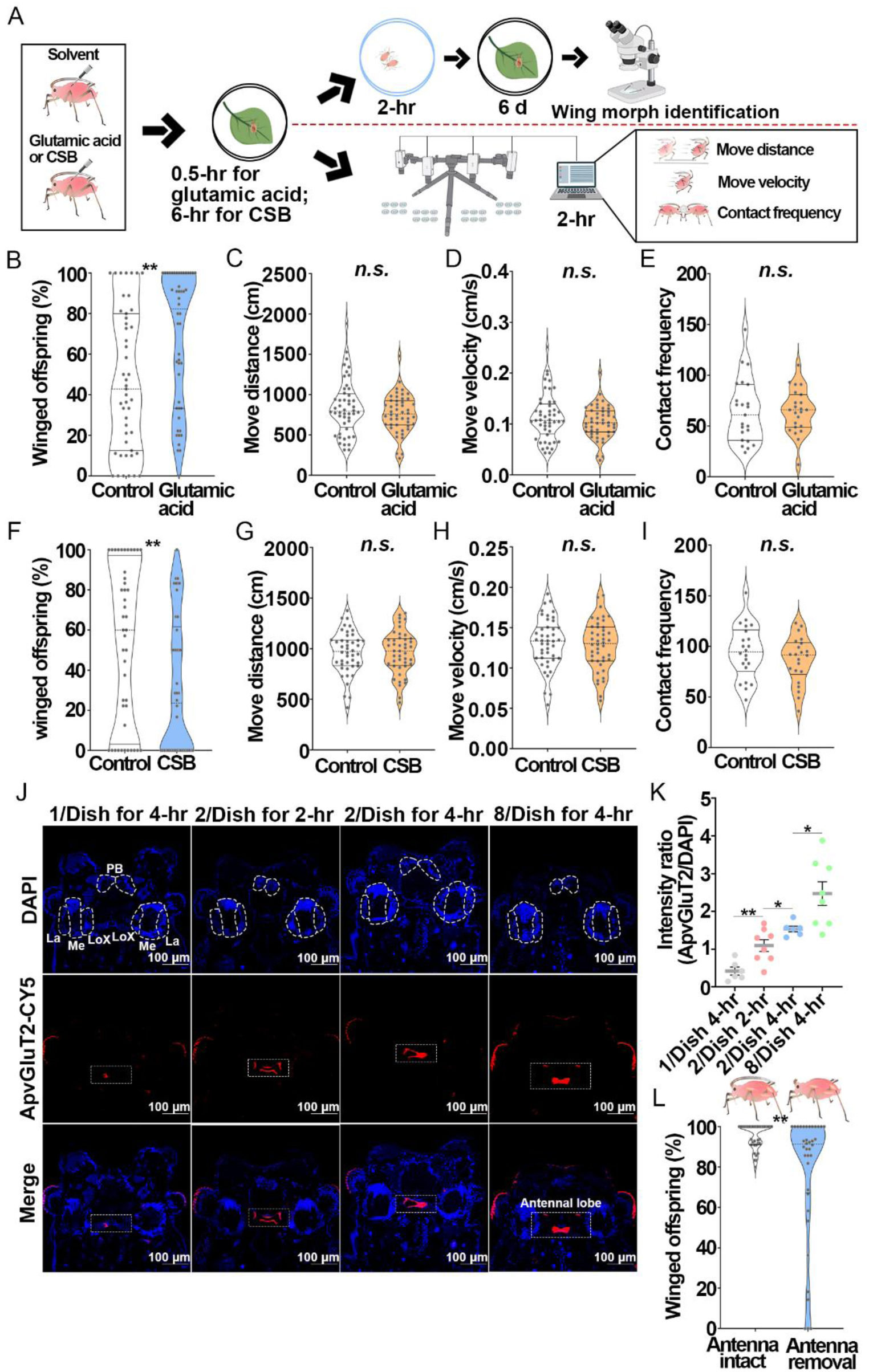
The effect of *ApvGluT2* on the proportion of winged offspring and its location in aphid brain. (A) Schematic diagram showing the procedure of winged offspring inspection and behavior monitoring in pharmacological experiments in pea aphids. (B) Increased percentage of winged offspring by glutamic acid injection, an agonist of vGluT (n>47). (C) The effect of glutamic acid injection on the movement distance (n=48), (D) the movement velocity (n = 48), and (E) the contact frequency of maternal aphids (n=24). The Kolmogorov-Smirnov test was used to compare means. (F) Decreased percentage of winged offspring by CSB injection, an antagonist of vGluT (n>42). (G) The effect of CSB injection on the movement distance (n=44), (H) the movement velocity (n=44), and (I) the contact frequency of maternal aphids (n=22). (J) mRNA-FISH. *ApvGluT2* was mainly expressed in the neural antennal lobes (highlighted by white squares) of aphid brain and was induced by high maternal density and contact treatments. Cy5-conjugated *ApvGluT2* probe was shown in red, and nuclei were stained with DAPI in blue. The selected neuropils (La, lamina; Me, medulla; LoX, lobula complex; PB, protocerebral bridge) were outlined by dotted lines. (K) The relative intensity of *ApvGluT2* in the antennal lobes was quantified by LAS X (n >6). (L) The proportion of winged offspring was declined by antennae amputation (n=36). The Mann Whitney non-parametric test was used to compare means of the proportions of winged offspring. Student *t* test was used to compare means of relative intensity of *ApvGluT2*. *n.s.*, not significant, *P < 0.05, **P < 0.01.

### Embryonic *ApmGluR3* relayed the key transgenerational signal involved with wing dimorphism

Because the embryonic period is a critical time point for determination of offspring wing morph, we hypothesized that *ApmGluR3* in the embryo was a key regulator downstream of *ApvGluT2* that mediated maternal-offspring communication. Application of mGluR antagonist LY341495 decreased the winged offspring by 33% (Figure 3A), whereas injection of mGluR agonist (2R, 4R)-4-aminopyrrolidine-2,4-dicarboxylate (APDC) increased the proportion of winged offspring by 40.4% (Figure 3B). Topical application of ds*ApmGluR3-RNA* knocked down the expression of *ApmGluR3* by 62.2% in the embryo, and the percentage of winged offspring dropped from 56.3% to 36.5% (Figure 3C-E). To establish a causal link between brain vGluT2 and embryo mGluR3, ds*ApvGluT2-RNA* and APDC were administered simultaneously (Figure 3F-G). Notably, reduction of winged offspring caused by the knockdown of *ApvGluT2* diminished upon application of APDC, the agonist of embryonic mGluR3 (Figure 3 F-G). FISH assays revealed abundant *ApmGluR3* in stage 20 embryos (Figure 3H). More importantly, the expression of *ApmGluR3* in the wing disc area of the embryo was up-regulated by density and contact treatments (Figure 3H-I). Presumably, ApmGluR3 functioned downstream of ApvGluT2 in the signaling pathway and mediated transgenerational wing dimorphism during embryonic development.

**Figure 3.**
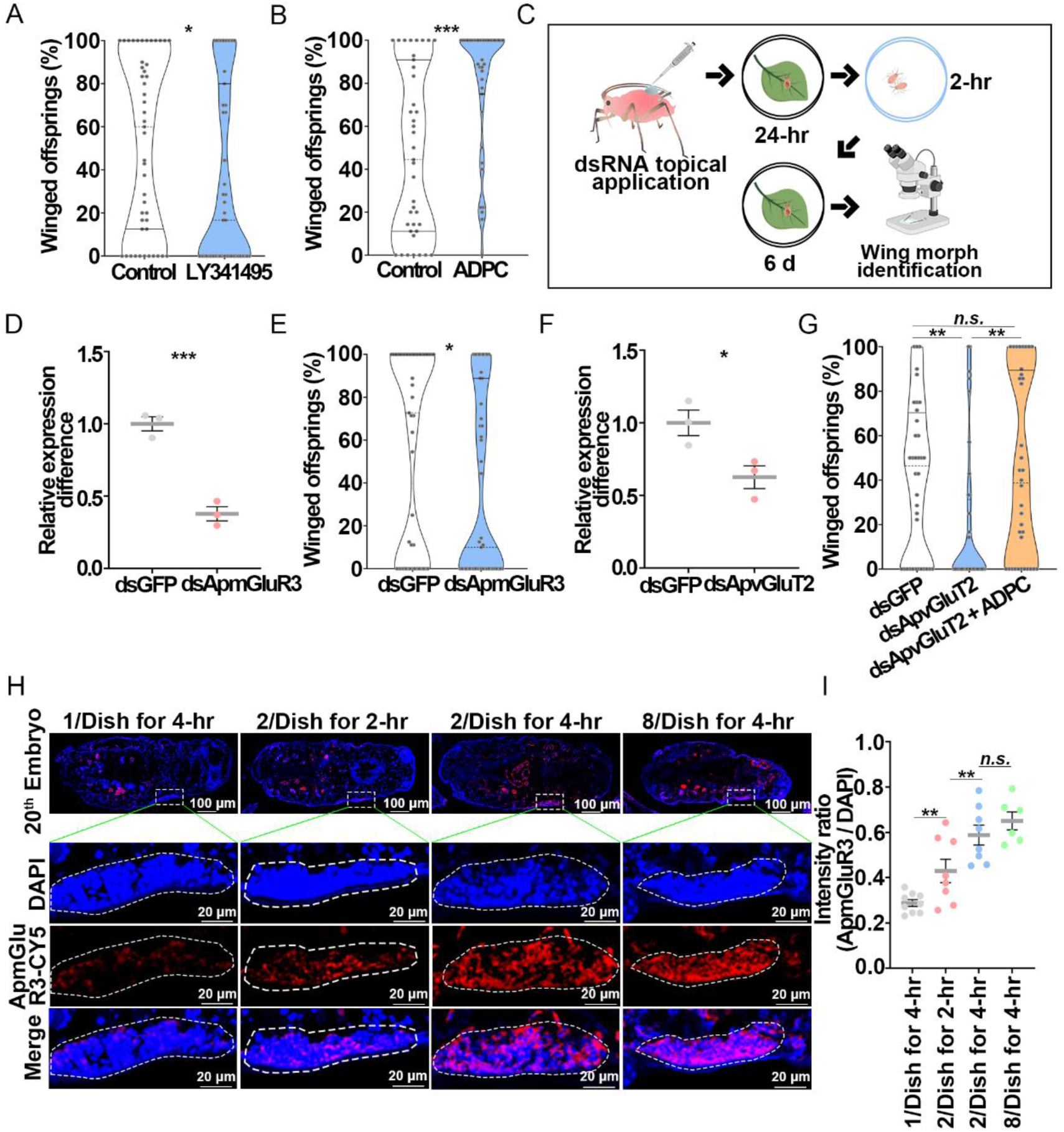
Embryonic *ApmGluR3* relayed the transgenerational signal from *ApvGluT2* about the induction of winged offspring. (A) Declined proportion of winged offspring by injection of an mGluR antagonist LY341495 (n=47), in contrast to (B) increased proportion by agonist ADPC (n=39). (C) RNAi experimental design. (D) Knockdown of *ApmGluR3* in the embryo (n=3), and (E) reduced proportion of winged offspring (n=47). (F) RNAi efficiency of *ApvGluT2* in the head (n=4). (G) Effect of ds*ApvGluT2* on the proportion of winged offspring was rescued by ADPC (n=36). (H) The expression of *ApmGluR3* in the stage 20 embryos increased as the contact duration increased. *ApmGluR3* probe conjugated with Cy5 was in red, and nuclei were stained with DAPI in blue. Wing discs shown in confocal were highlighted by white circles. (I) The relative intensity of *ApmGluR3* in the wing discs of stage 20 embryos was quantified by LAS X (n>9). Mann Whitney non-parametric test was used to compare the means of the proportion of winged offspring. Student *t* test was used to compare means of transcript level or the relative intensity of *ApmGluR3*. *P < 0.05, **P < 0.01, ***P < 0.001.

### ApmGluR3 promoted phosphorylation of ApFoxO in embryos

Our previous study indicated that the embryonic transcription factor, ApFoxO is a crucial suppressor of wing development in pea aphids (*Yuan et al., 2023a*). In planthoppers, phosphorylation of FoxO in wing bud cells caused its translocation from the nucleus to cytoplasm, thereby losing its transcriptional repressor function in wing development (*Lin et al., 2016*; *Xu et al., 2015*). We therefore examined whether ApmGluR3 affected phosphorylation of *Ap*FoxO in embryos (Figure 4A). Initially, we confirmed that ApFoxO indeed suppressed winged offspring: knockdown of *ApFoxO* in embryos of maternal aphids by 28.6% (*P* value < 0.0001; Figure 4B) led to a significant increase in the percentage of winged offspring, from 53.9% to 61.8% (*P* value = 0.0255; Figure 4C). Amino acid sequence alignment of ApFoxO with its homologs in fruit fly (DmFoxO) and human (hFoxO4) identified three conserved putative phosphorylation sites in ApFoxO, T15, S187, and S250 (Figure S3). Akt could further phosphorylate these sites to sequestrate FoxO in the cytoplasm, thus preventing FoxO factors from transactivating their target genes (*Calnan and Brunet, 2008*). Our results found that the phosphorylation of ApFoxO was successfully detected by antibody against phosphorylated T15, but failed to be detected with antibodies targeting phosphorylated-S187 and S250 (Figure 4D-F, S4). Physical contact for 4-hr significantly increased phosphorylated *Ap*FoxO in embryos (by 59.8%) compared to the solitary counterparts (Figure 4D). Knockdown of *ApmGluR3* or *ApvGluT2* reduced the phosphorylation level of ApFoxO in embryos by 21.3% and 54.3%, respectively (Figure 4E-F). In addition, applying the specific Akt inhibitor MK-2206 remarkably reduced the percentage of winged offspring, from 49.8% to 24.9% (Figure 4G). These results implied that the ApvGluT2-ApmGluR3 cascade increased the phosphorylation of ApFoxO in embryos, promoting the winged offspring.

**Figure 4.**
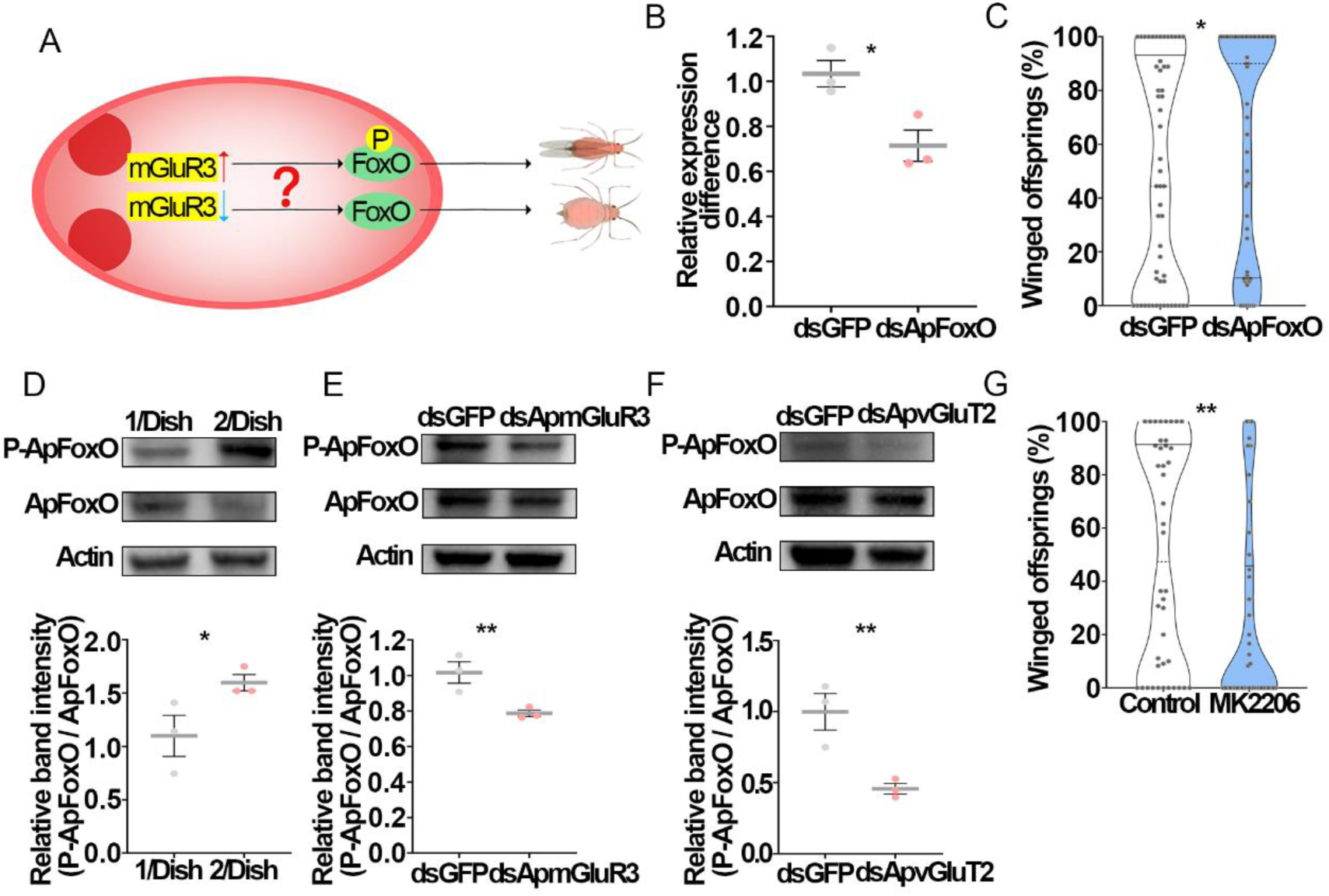
ApvGluT2 and ApmGluR3 promoted phosphorylation of embryonic ApFoxO, facilitating the formation of winged aphids. (A) Hypothetical model of *Ap*mGluR3-regulated phosphorylation of *Ap*FoxO in the embryo that controls the wing dimorphism of aphids. (B) RNAi of *ApFoxO* in embryos (n>5), and (C) its effect on the proportion of winged offspring (n>54). (D) The phosphorylated ApFoxO level in the embryo was enhanced by two-adult contacting for 4-hr treatment(n=3) and (E) reduced by ds*ApmGluR3*, and (F) ds*ApvGluT2* (n=3), as determined by western blotting analyses. (G) The proportion of winged offspring was reduced by Akt antagonist MK2206 that directly targeted FoxO as a substrate (n=42). Mann Whitney non-parametric test was used to compare the means of the proportion of winged offspring. Student *t* test was used to compare means of gene transcript or protein level. *P < 0.05, **P < 0.01, ***P < 0.001.

### ApFoxO bound to *ApHh* promotor to suppress its activation

Our previous study found that the transcripts of four wing development genes, including *Hedgehog* (*Hh*), *Vestigial* (*Vg*), *Decapentaplegic* (*Dpp*) and *Mob as tumor suppressor* (*Mats*), were significantly altered by knockdown of *ApFoxO* in the embryos of pea aphids (*Yuan et al., 2023a*). The stage 20 embryo appears to be a critical point in development for an aphid to determine irreversibly its winged or wingless morph (*Ishikawa and Miura, 2013*; *Ogawa and Miura, 2013*). Knockdown of *ApmGluR3* at stage 20 embryo decreased expression of *ApHh* and *ApMats* by 37.7% and 41.5% (Figure 5A). Interestingly, knockdown of *ApFoxO* caused an increase in *ApHh* transcripts, but a decrease in *ApMats* in stage 20 embryos (*Yuan et al., 2023a*). Accordingly, we ruled out *ApMats* and focused on *ApHh* as the suppressing target of ApFoxO. The application of ds*ApHh-RNA* reduced the expression of *ApHh* by 62.4% in stage 20 embryos, and decreased the proportion of winged offspring from 42.5% to 23.5% (Figure 5B-C). Knockdown of *ApFoxO* in stage 20 embryo increased the proportion of winged offspring by 40.4%. Percentage of winged offspring returned to the control level when both *ApFoxO* and *ApHh* were silenced (Figure 5D-E). These findings demonstrated that ApFoxO repressed wing development via inhibiting *ApHh* expression at the 20^th^ embryo stage.

**Figure 5.**
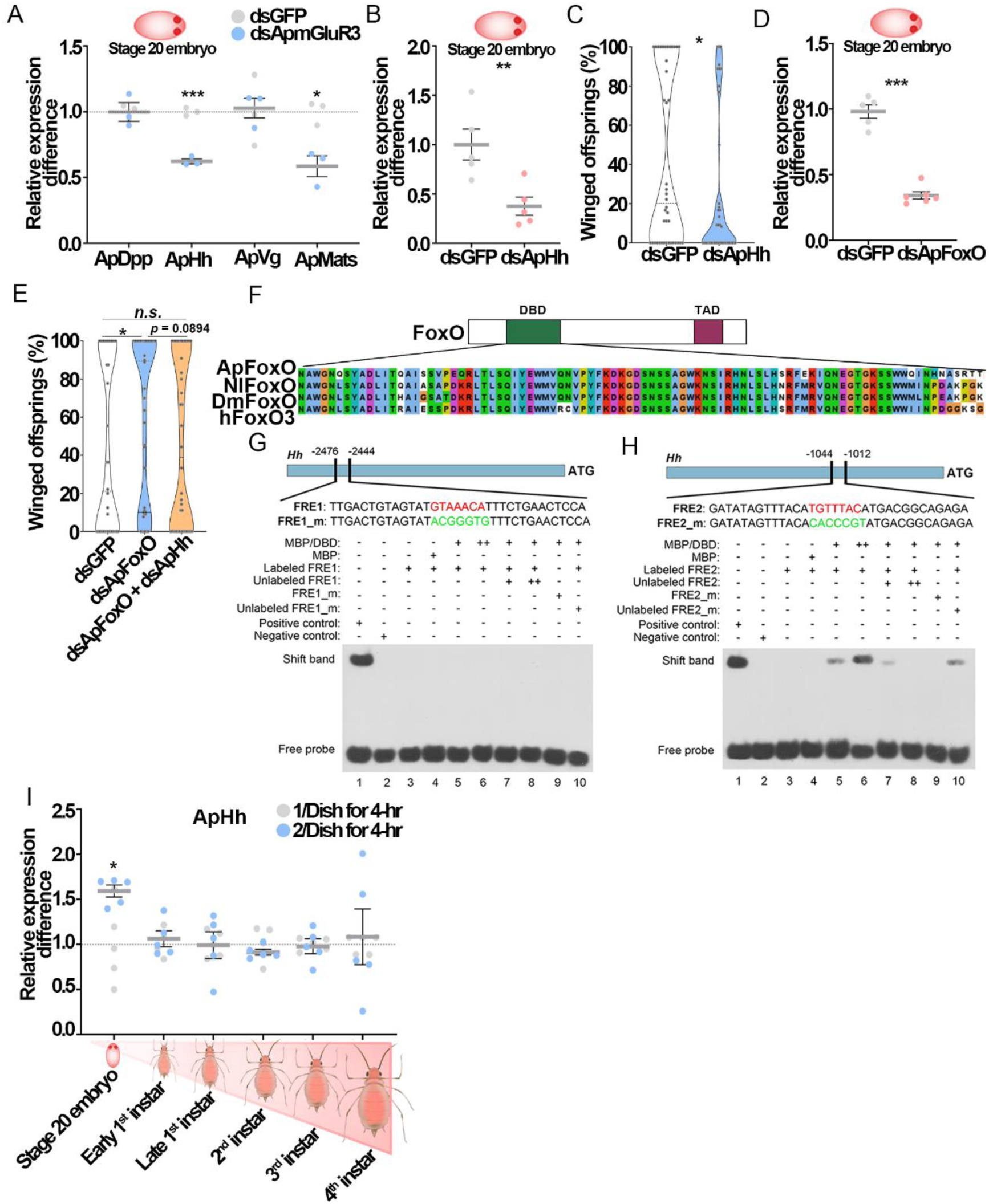
ApFoxO transcriptionally suppressed *ApHh* at stage 20 embryo. (A) Knockdown of *ApmGluR3* significantly reduced the transcripts of *hedgehog* (*ApHh*) and *Mob as tumor suppressor* (*ApMats*) in stage 20 embryos (n=3). (B) RNAi efficiency of *ApHh* in stage 20 embryos (n=5). (C) The effects of ds*ApHh* on the proportion of winged offspring (n=42). (D) RNAi efficiency of *ApFoxO* in stage 20 embryos (n=5). (E) The proportion of winged offspring was rescued by knocking down of *ApHh* in ds*ApFoxO* aphids (n>40). (F) Sequence alignment of DNA binding domains (DBD) among FoxO homologs from brown planthopper (NlFoxO), *Drosophila* (DmFoxO) and human (hFoxO). (G) EMSA showing that Maltose-binding protein (MBP)/ DNA-binding domain (DBD) of ApFoxO fusion failed to bind to *FRE1* (FoxO recognition element 1), but (H) bound to *FRE2* on the *ApHh* promoter. Lane 1: positive control. Lane 2: negative control. Lane 3: biotin-labeled *FRE* only. Lane 4: 0.5 μg recombinant MBP +labeled FRE. Lane 5: 0.5 μg or Lane 6: 5 μg of recombinant MBP/DBD fusion protein + labeled *FRE*. Lane 7: 0.5 μg recombinant MBP/DBD fusion protein + labeled *FRE* + 20 μmol (5 ×) unlabeled *FRE* competitor, or Lane 8: 25 × unlabeled competitor. Lane 9: 0.5 μg recombinant MBP/DBD fusion protein + 5 × biotin-labeled mutant FRE (FRE_m). Lane 10: 0.5 μg recombinant MBP/DBD fusion protein + biotin-labeled FRE + 5 × biotin end-labeled FRE_m. Biotin-labeled *FRE* was all 4 μmol in each lane. (I) Two-adult contacting increased *ApHh* transcripts only at stage 20 embryos relative to solitary aphids (n=5). Mann Whitney non-parametric test was used to compare the means of the proportion of winged offspring. Student *t* test was used to compare means of gene transcript level. *n.s.* not significant, *P < 0.05, **P < 0.01, ***P < 0.001.

Electrophoretic mobility shift assays (EMSA) were then performed to detect binding of ApFoxO to the promotor of *ApHh* via the FoxO recognition element (*FRE*), TTGTTTAC. Typically, FoxO interacts with the *FRE*-element through its highly conserved DNA-binding domain (DBD) consisting of ∼100 amino acid residues to regulate transcription of target genes (*Furuyama et al., 2000*; *Xuan and Zhang, 2005*). The sequence alignment showed that DBD of *ApFoxO* shared high sequence similarity to its homologs in brown planthopper *Nilaparvata lugens*, fruit fly *Drosophila melanogaster* and human *Homo sapiens* (Figure 5F). Two FREs, FRE1 and FRE2, were identified in the 3-kb segment upstream of the transcription start site of *ApHh* (Figure 5G-H). Recombinant maltose-binding protein (MBP)-DBD_ApFoxO_ was expressed in *Escherichia coli* and purified for EMSA (Figure S5). Biotin-labeled probes of FRE2 result in mobility shift in a concentration-dependent manner, while no binding to biotin-labeled FRE1 was detected (Figure 5G-H). Notably, the intensity of shift bands was greatly reduced by adding the unlabeled FRE2 probe. No binding occurred when the cis-element was mutated (Figure 5H), supporting specific interaction between *FRE2* and ApFoxO.

We further compared the temporal expression patterns of *ApHh* between offspring collected from solitarious and the 4-hr physical contact treatment across the developmental stages from stage 20 embryos to the 4^th^ instar nymphs. Little treatment difference was observed in *ApHh* transcripts in postnatal development stages, suggesting that the stage 20 embryo was a critical time point for decision-making regarding the developmental trajectory of the wing disc (Figure 5I). Together, these findings indicated that embryonic ApFoxO and ApHh were the executors to determine winged dimorphism in aphids.

## Discussion

The phenomenon where a single genotype is expressed in a spectrum of phenotypes, both within a single generation and across successive ones, is central to comprehension of how populations adapt to their environments. This adaptability is particularly complicated in the context of TPP due to its obscure nature of signal transduction mechanisms facilitating parent-offspring communication. Although some understanding has been attained about within-generation wing plasticity in insects, much less is known regarding the transgenerational regulatory mechanism. Our research has demonstrated that maternal brain *ApvGluT2*, mainly expressed in antennal lobes, and embryonic *ApmGluR3* were specifically up-regulated when aphids experienced physical contact or high population density. This was followed by an elevated phosphorylation of *Ap*FoxO in stage 20 embryos, which caused the loss of its suppression of *ApHh*, as well as a high proportion of winged offspring (Figure 6). Conversely, lower expression of *ApvGluT2* and *ApmGluR3* in solitary aphids sustained a higher level of unphosphorylated ApFoxO, which effectively suppressed *ApHh* expression and favored the production of wingless offspring. These findings showcased how pea aphids deployed the glutamate signaling pathway in regulating transgenerational wing dimorphism in response to environmental cues, and highlighted the importance of Hh as secreted protein governing wing development trajectory in stage 20 embryos.

**Figure 6.**
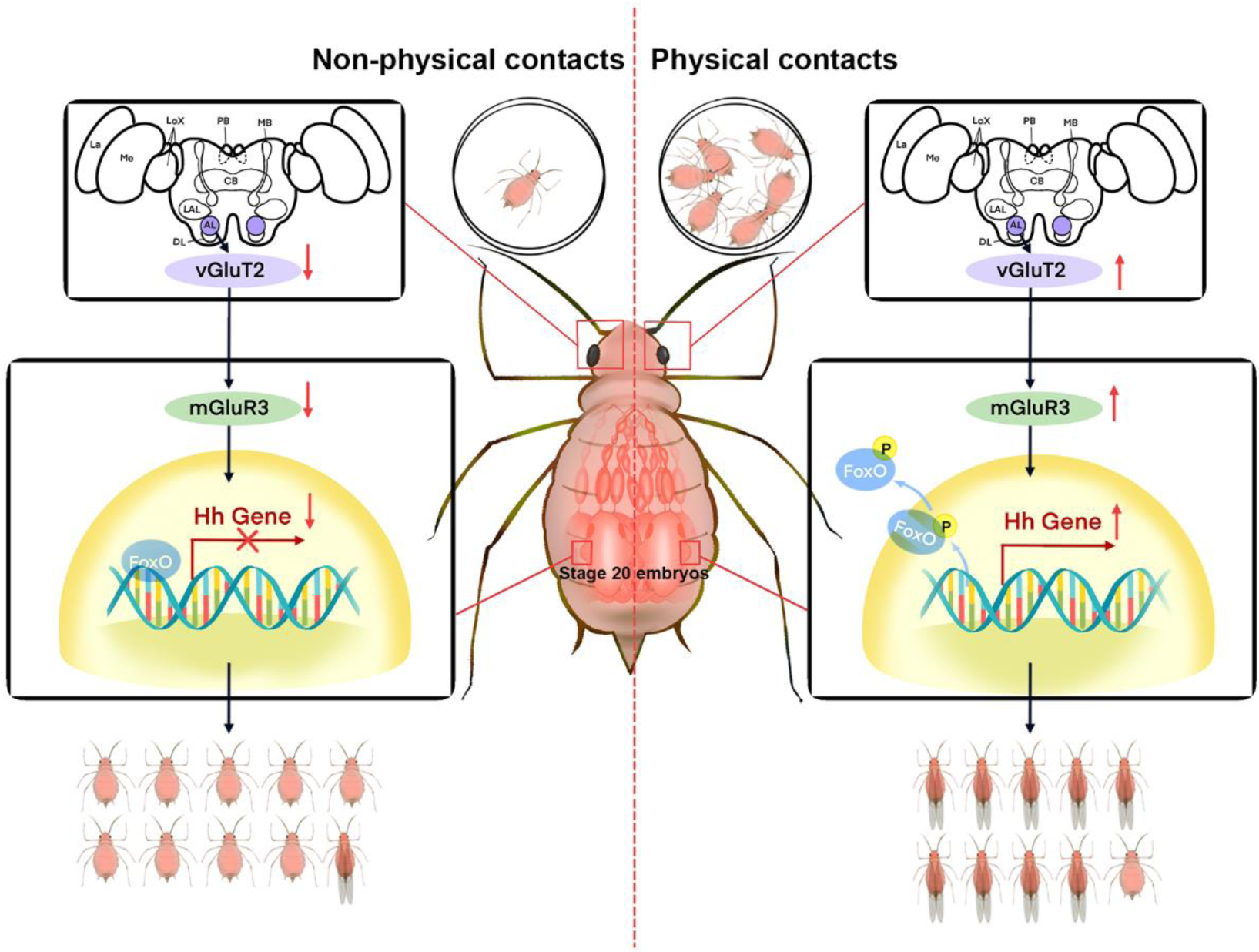
Illustration of how the ApvGluT2-mGluR3 module transgenerational controls wing morphs in response to physical contact. In solitary aphids, relatively high level of unphosphorylated ApFoxO binds to the *cis*-element of *ApHh* promoter, suppressing its expression at stage 20 embryo. As a result, the prevalence of wingless offspring was observed. Once experienced physical contact, maternal aphids increased the *ApvGluT2* expression in antennal lobes of the brain and *Ap*m*GluR3* in wing discs of the embryo. This is followed by increased phosphorylation of ApFoxO. The phosphorylated ApFoxO is transported from the nucleus to the cytoplasm, thereby releasing the transcriptional suppression of *ApHh*, leading to high proportion of wingless offspring. vGluT2, vesicular glutamate transporter 2; mGluR3, metabotropic glutamate receptor 3; FoxO, Forkhead transcription factor subgroup O; Hh, hedgehog; La, lamina; Me, medulla; Lox, lobula complex; PB, protocerebral bridge; MB, mushroom body; CB, central body; LAL, lateral accessory lobe; AL, antennal lobe; DL, dorsal lobe.

High population density is a reliable environmental cue for intense competition or deteriorating habitat conditions for individuals, resulting in a high proportion of winged offspring. The tactile, olfactory, and visual signals are essential cues for aphids to perceive population density and trigger appropriate phenotypic changes (*Topaz et al., 2012*). For instance, physical contact could induce gregarious locusts from a solitary population by releasing aggregation pheromone 4-vinylanisole (*Guo et al., 2020*; *Yang et al., 2023*). In aphids, the increased frequency of physical contact due to increasing population density could also explain wing induction (*Sutherland, 1969*). A “pseudo-crowding” hypothesis has been proposed to explain the predator-induced wing induction in aphids. The perception of predators or the alarm pheromone increases the locomotion activity of aphids, causing higher frequency of physical contact among individuals, similar to what happens when aphids are crowded. Although single individuals could behaviorally react to the alarm pheromone, lack of physical contact renders them unable to trigger wing production (*Kunert et al., 2005*). Consistently, aphids with ablated antennae fail to produce more winged offspring in the presence of predators (*Kunert et al., 2005*). Our previous screening of natural pea aphids gave rise to a strain that was highly sensitive to physical contact in triggering winged offspring, which provided an excellent biotype for dissecting the underlying molecular mechanism (*Yuan et al., 2023b*). Our results found a strong correlation between the locomotor activity of maternal aphids and the proportion of winged offspring in the treatment of two-adult contacting for 2-hr. Presumably, elevated mobility of maternal aphids led to higher frequency of physical contact, in agreement with the pseudo-crowding hypothesis.

Low titers of serotonin, dopamine and octopamine are found in densely populated pea aphids (*Vellichirammal et al., 2016*). Of these biogenic amines, only dopamine was reported to be a suppressor for winged morph production (*Liu and Brisson, 2023*). In this study, physical contact stimulated higher transcription activity of *ApvGluT2* mainly located in the antennal lobes of the maternal brain. Blocking the vGluT2 activity reduced the proportion of winged offspring but did not alter maternal locomotor activity. It has been shown that vGluTs are essential in sensing mechanical pain in somatosensory system, dorsal horn and brain (*Seal, 2016*). As evidenced by transiently expressing *vGluT3* in neurons of mammal deep dorsal horn, it could directly receive Aβ primary sensory input caused by mechanical stimuli, and relay polysynaptic transmission onto lamina I pain processing neurons (*Peirs et al., 2015*; *Seal et al., 2009*). Our phylogenetic analysis and sequence alignment indicated that ApvGluT2 belonged to animal Clade III in the solute carrier (SLC) 17 subfamily, sharing 21.33%, 22.33%, and 20.74% amino acid identity with human vGluT1, vGluT2, and vGluT3, respectively, along with 12 conserved transmembrane regions (Figure S6 A-B). Notably, ApvGluT2 exhibited a significant structural similarity to human vGluT2, as evidenced by minimal root mean square deviation (RMSD) value of 4.131 Å, suggesting ApvGluT2 maintained relatively conserved function in aphids (Figure S6 C). Most likely, a specific group of *ApvGluT2*-expressing neurons in antennal lobes of aphids received physical contact-induced sensory input, then transmitted the synaptic signals to downstream nerves to modulate the offspring wing morph at their embryonic period.

Considering that the antennal lobes are located at a considerable distance from the embryos, an important issue is how the brain ApvGluT2 controls the wing development trajectory in embryos. Our results revealed that the embryonic *ApmGluR3* was up-regulated by maternal high density and physical contact experiences. Although *ApmGluR3* was expressed throughout the embryo, it could be up-regulated by density and contact treatments in the area of wing discs. Injection of mGluR agonist rescued the negative effect of ds*ApvGluT2* on the proportion of winged offspring, suggesting *ApvGluT2* and *ApmGluR3* coordinately regulated the production of wing morph in offspring triggered by maternal density. Early studies have shown that the vGluT-mGluR signal axis orchestrates the function of specific neuronal networks involved in motor coordination, emotions, and cognition (*Ibrahim et al., 2020*). Furthermore, the glutamatergic neurons in the ventral nerve cord and thoracic abdominal ganglion could project to the reproductive tract to regulate reproduction (*Castellanos et al., 2013*; *Deshpande et al., 2022*; *Rodríguez-Valentín et al., 2006*). Therefore, it is likely that glutamatergic neurons in antennal lobes of aphids recruited vGluT2-mediated signaling to activate the glutamatergic neurons in the ventral nerve cord and thoracic abdominal ganglion, innervating the mGluR3-expressed cells in the wing disc of embryos to control their wing development.

Although FoxO acts as a major suppressor of the winged morph in both within- and trans-generational wing polyphenism, the regulatory routes may vary in developmental stages by targeting different downstream wing development genes. Some such as *Vg* and *Ubx* that control the wing patterning in wing buds could be suppressed by FoxO within a generation (*Liu et al., 2020*; *Zhang et al., 2021*). As for the transgenerational regulation, we showed that physical contact-induced increases of brain *ApvGluT2* and embryonic *ApmGluR3* enhanced the phosphorylation level of ApFoxO, as well as the proportion of winged offspring. In contrast, knockdown of *ApvGluT2* and *ApmGluR3*, and the application of Akt antagonist could reduce the phosphorylation level of ApFoxO, consistent with a previous study, where mGluR activates PI3K in *Drosophila* larval motor neuron to alter phosphorylation of FoxO (*Howlett et al., 2008*).

During the differentiation of Drosophila imaginal disc, *Hh* is transcribed in the posterior compartment of the wing disc and acts as a short-range morphogen that is secreted anteriorly, establishing a concentration gradient that provides positional information to wing disc cells and determines the developmental trajectory of the wing (*Hartl and Scott, 2014*; *Neto-Silva et al., 2009*). *Hh* controls the expression of the long-range morphogen *Dpp* to regulate the pattern and growth of the wing disc through its essential downstream component *Cubitus interruptus* (*Bejarano et al., 2007*). In this study, the embryonic expression of *ApDpp* was not significant for ds*GFP* vs. ds*ApmGluR3* group and solitary vs. physical contact group, leaving the possibility that the clean sampling of wing discs of stage 20 embryos instead of the whole body could reveal the wing gene regulatory network (GRN) in pea aphids. Moreover, knockdown of *ApmGluR3* did not affect *vestigial*, which was wing specific and necessary for wing development in insects, suggesting *ApmGluR3* indirectly regulated the wing GRN in embryos, and the connection between *ApmGluR3* and the wing GRN could be mediated by other downstream developmental or physiological processes that affected wing morph development. Furthermore, our data demonstrated that FoxO could directly bind to the promotor of *ApHh* to suppress its expression in stage 20 embryos. Beyond the role in wing development, Hh functions as a secreted signaling protein in many other organs, regulating a range of processes including embryonic development, tissue homeostasis, organ regeneration, axon guidance, and cellular metabolism (*Briscoe and Thérond, 2013*). Since the wingless and winged aphids exhibit a range of differences in morphological, behavioral and life history (*Brisson, 2010*), the different expression of Hh in stage 20 embryos could be critical to aphid wing-morph differentiation in response to high population density. Together, we have provided a novel mechanistic insight into the regulation of TPP and demonstrated, for the first time that the glutamate signal transduction cascade mediates the transgenerational wing dimorphism in aphids.

## Materials and methods

### Aphid rearing

The pea aphid *A. pisum* (strain: Ningxia Red) was originally collected from *Medicago sativa* in Ningxia Province and had been reared in the laboratory for over 8 years. Nymphs from the same parthenogenetic pea aphid female were reared on *Vicia faba* at 18-20°C, with 60% relative humidity and a photoperiod of a 16 hr: 8 hr (light/dark) cycle. To eliminate the transgenerational effects on offspring morphs, females were maintained at low density (three per plant) on *V. faba* seedlings for more than three generations (*Sutherland, 1969*). This strain was used to test the effects of two-adult physical contact or crowding on the proportion of winged offspring.

### Induction of winged offspring

To effectively induce a high proportion of winged offspring, the maternal density and contacting duration were evaluated in petri dishes (35 mm in diameter) as described by 14. Three groups of aphids consisting of 1, 2, or 8 wingless adults, respectively, were placed in a petri dish for 0.5-, 2-, and 4-hr. After treatment, each adult was transferred to a freshly detached *V. faba* leaf kept in petri dishes with 1% agar to reproduce for 24-hr. After newborns reached the fourth instar, the proportion of winged offspring from each adult was calculated.

For density treatment, two groups of 1 and 15 maternal aphids were placed on fava seedlings for 24-hr. This was followed by transferring each adult to a fresh *V. faba* leaf as described above.

### Aphid locomotor activity tracking in petri dishes

One or two maternal aphids were placed in a petri dish (diameter 35 mm), and locomotor activities of each aphid within 2- or 4-hr were recorded (25 frames/s) with a video camera. Six or 12 aphids in each treatment were simultaneously recorded per replicate. The valid video data for each treatment were collected and analyzed at least 40 replicates. Videos were analyzed using the EthoVision XT software (v.11.5, Noldus Information Technology) to measure the total distance (moving distance, unit: cm) and the average velocity of movement (move velocity, unit: cm/s), as well as the contact frequency. Physical contact between two maternal aphids was considered to have occurred when their proximity distance was less than 0.5 cm.

### RNA-Seq

Maternal aphids housed individually or two per petri dish for 4-hr were used for differential gene expression analyses. The heads and embryos were dissected from 10 and 5 adult aphids per replicate, respectively. Four of such biological replicates were collected RNA-seq. DEGs were identified with DESeq2 R package (1.20.0) using the expected number of Fragments Per Kilobase of transcript sequence per Million base pairs sequenced. The sequencing data were deposited in the Short Read Archive (NCBI) under accession number PRJNA1017088.

### RT-qPCR

Total RNA was extracted using TRIzol reagent (Invitrogen, CA, USA), and reverse transcribed using the PrimeScriptTM RT reagent Kit with gDNA Eraser (Perfect Real Time) kit (Takara, Dalian, China). The RT-qPCR reactions were carried out on the PikoReal 96 Real-Time PCR System (Thermo) using the SuperReal PreMix Color (SYBR Green) kit (Tiangen, Beijing, China). Three technical replicates were applied for each biological replicate. The housekeeping gene ribosomal protein L27 (RPL27) was used as the internal qPCR standard.Gene expression data were analyzed by relative quantification with the 2^-△△CT^ method (*Livak and Schmittgen, 2001*). Specific primers for each gene were designed using Primer Premier 6 and tested in preliminary experiments. The oligonucleotides are listed in Table S1.

### Pharmacological experiments

Glutamate signaling agonist glutamic acid (Topscience, Shanghai) and mGluR agonist (2R, 4R)-ADPC (Enzo Life Sciences, New York) were dissolved in water at concentrations of 33.98 mM and 100 nM, respectively. vGluT2 antagonist Chicago Sky Blue 6B (CSB) (Topscience, Shanghai), mGluR antagonist LY341495 (MedChemExpress, New Jersey), and Akt inhibitor MK-2206 (Selleckchem, Houston) were dissolved in DMSO at concentrations of 10 mM, 16.98 mM, and 5 mM, respectively. A volume of 100 nL of each chemical, except for MK2206, which was 23.6 nL, was delivered into hemolymph from dorsal abdomens using the Nanoject II microinjection system (Drummond Scientific Company, Broomall, PA, USA). An equivalent amount of solvent was used as a solvent control.

### Fluorescence in situ hybridization

FISH was performed with the technique modified by Kliot et al. (*2014*). After washing in TBST (TBS with 0.2% Triton-X) for 10 min, sample section slices were rinsed three times in the hybridization buffer (20 mM Tris-HCl, pH 8.0, 0.9 M NaCl, 0.01% sodium dodecyl sulfate, and 30% formamide), and hybridized overnight in the hybridization buffer containing 10 pmol of the fluorescent RNA probe (conjugated with FAM or Cy5). After 3x rinses in TBST, samples were mounted in Fluoroshield Mounting Medium with DAPI (Abcam). Sections were imaged using a Leica M205C confocal microscope (Zeiss, Germany). The fluorescence intensities were quantified using Leica Application Suite (LAS) software.

### RNA interference

Double-stranded RNA (dsRNAs) i.e. ds*ApvGluT2*, ds*ApmGluR3*, ds*ApFoxO*, ds*ApHh*, and the ds*GFP* control were synthesized by T7 RiboMAX Express RNAi System (Promega, P1700) according to the manufacturer’s protocol. Each (2 μg/μl) was mixed with the nanocarrier in a water solution at a volume ratio of 1:1. A volume of 1 μL of the solution containing 1 μg nanocarrier/dsRNA complex was topically applied to the dorsal side of the female abdomen. Embryos from aphids or heads from ten aphids in each sample were collected 24 h after topical application of dsRNA for testing silencing efficiency using qPCR. Each treatment contained at least three biological replicates. The primers used for dsRNA synthesis are listed in Table S2.

### Antibody preparation and western blotting analysis

Three polyclonal rabbit antibodies against phosphorylated T15, S187 and S250 of ApFoxO were prepared by ABclonal using synthetic peptides RARSNT(p)WPLPR (10 to 20 aa), RRRAVS(p)METPK (182 to 192 aa) and RARASS(p)NASS (245 to 254 aa) as the immunogens. Western blotting was performed to detect phosphorylation levels of FoxO in embryos of adult females. The antigen–antibody complexes were visualized using a secondary goat anti-rabbit IgG (H+L)-conjugated horseradish peroxidase (HRP) antibody (LABLEAD).

### Electrophoretic mobility shift assays (EMSA)

The FoxO DNA-binding domain (DBD) (89 to 178 aa) was cloned into the pMAL-c2x expression vector, which contained an N-terminal maltose-binding protein (MBP), and subsequently expressed in *E. coli* strain Rosetta (DE3). Proteins were purified from cell lysates using the pMAL protein fusion and purification system (New England Biolabs). The MBP protein was also independently expressed in pMAL-c2x as control. EMSA was conducted as described previously (*Hellman and Fried, 2007*). Probes containing the DBD binding sites were labeled with biotin at the 5’ end. Unlabeled probe was used as the competitor. The probes were incubated with purified MBP/DBD recombinant protein at room temperature for 30 min, run on a non-denaturing 0.5×TBE 6% polyacrylamide gel for 1h (60 V at 4°C), and transferred onto Biodyne® B nylon membrane (Pall Corporation). Signals were visualized using the ChemiDoc XRS kit (Bio-Rad Laboratories, UAS).

### Phylogenetic construction of SLC17 proteins

Protein sequences, 333 from 10 different species, were obtained from UniProt (https://www.uniprot.org/). Sequences were aligned with MAFFT v7.310 (*Katoh et al., 2002*) and automated trimmed with trimAl v1.4 (*Capella-Gutiérrez et al., 2009*). Phylogenetic analysis was performed using IQ-TREE v2.0.3 with LG+R9 model and ultrafast bootstrap approximation with 1000 replicates (*Minh et al., 2020*). Phylogenetic tree was visualized in Interactive Tree Of Life (iTOL) (*Letunic and Bork, 2021*).

### Protein Sequence and Structure analysis

The multiple sequence alignment was performed using Multiple Protein Sequence Alignment (MUSCLE). High confidence protein structure of *Ap*vGluT2 was predicted by Alphafold v2.3.0 (*Jumper et al., 2021*). The pairwise structural alignments were performed using PyMOL 2.6.0a0 (www.pymol.org).

### Statistical Analyses

The GraphPad software was used for statistical analyses. All data were checked for normality by the Wilk–Shapiro test. Two-tailed unpaired t-test was used to separate the means of normally distributed data, while Mann–Whitney test was used to analyze nonparametric data. All western blotting and confocal assays were repeated independently at least three times with similar results. Pearson’s correlations were calculated to determine the relationship between proportion of winged offspring and maternal locomotor activities.

## Data Availability

All data used in the study are included, and some are presented as supplementary information. All protocols have been described in Materials and Methods or in the references therein. No custom codes or mathematical algorithms were used in this manuscript.

## Author contribution

Yiyang Yuan project designing, performing experiments, data analysis and manuscript writing; Yanyan Wang, Wanwan Ye, Liqiang Xie, Fang Dong performing experiments; Keyan Zhu-Salzman manuscript writing and revising, and Erliang Yuan, Huijuan Guo, Shifan Wang manuscript revising; Feng Ge, Yucheng Sun project designing and coordination, manuscript writing.

## Declare of interests

We declare no competing interest.

## Acknowledgements

We thank Prof. Jinfeng Chen for his support in phylogenetic analysis. This project was supported by the National Key R&D Program of China (no. 2023YFD1400800), the Taishan Scholars Program, the Introducing Top Talent Program of Shandong (2023YSYY-006), the Agricultural Science and Technology Innovation Project of the Shandong Academy of Agricultural Sciences (no. CXGC2023F04), the National Natural Science Foundation of China (nos. 31970453, 32250002 and 32372636), and the State Key Laboratory of Integrated Management of Pest Insects and Rodents (no. IPM2306).

