## Supplemental Figure S1 for "The maternal vGluT2 and embryonic mGluR3 signaling relay system controls offspring wing dimorphism in pea aphid"

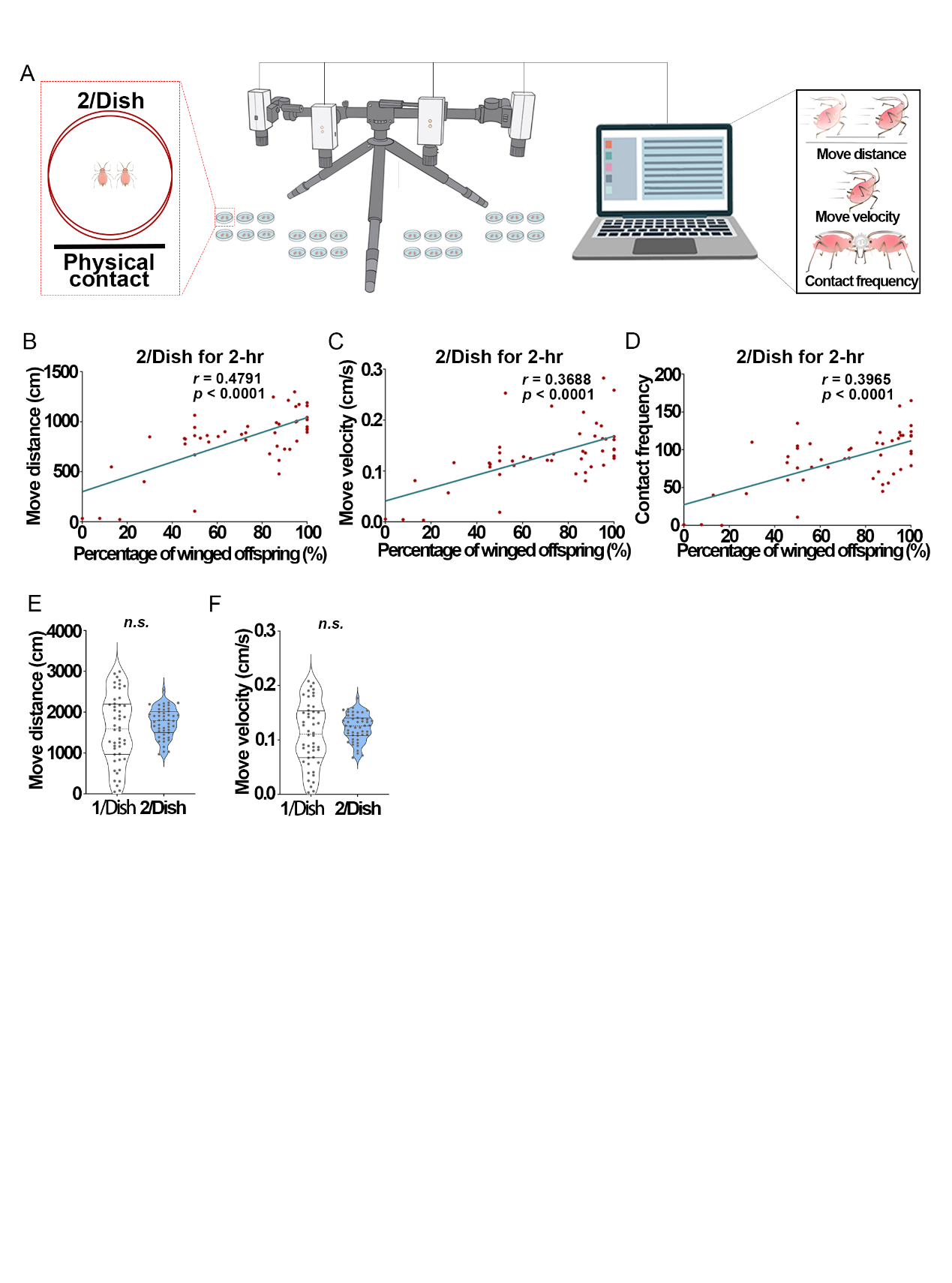

**Figure S1 Correlation between the proportion of winged offspring and maternal locomotor activity.** (A) Schematic diagram of locomotor activity monitoring. (B) Linear regression of the proportion of winged offspring vs the move distance, or (C) the move velocity, or (D) the contact frequency in maternal aphids. (E) The movement distance and (F) velocity of maternal aphids subjected to the two-adult contacting for 4-hr were similar to those of solitary maternal aphids. Student *t* test was used to compare means, n = 48, *n.s.* not significant.
