## Supplemental Figure S2 for "The maternal vGluT2 and embryonic mGluR3 signaling relay system controls offspring wing dimorphism in pea aphid"

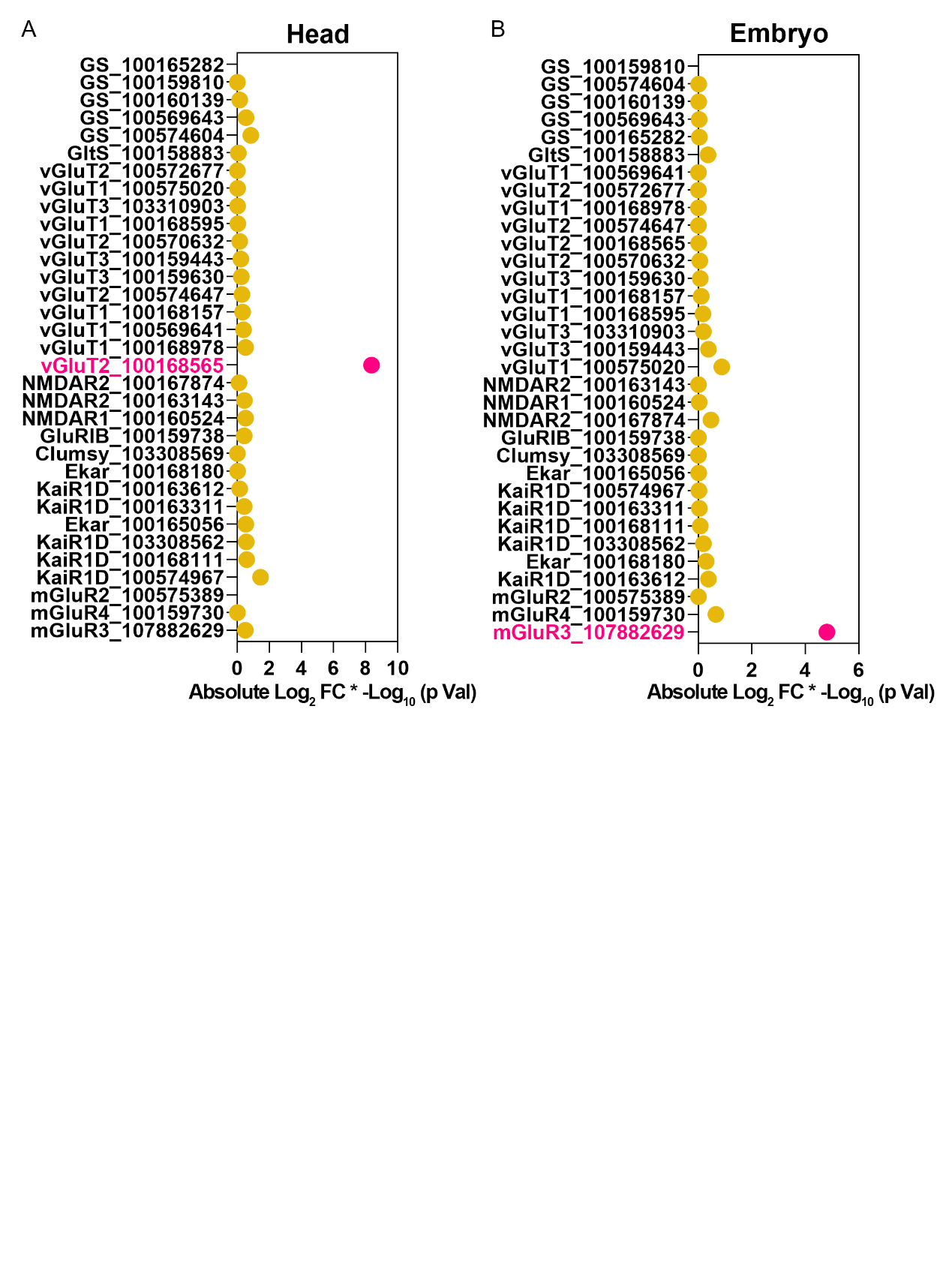

**Figure S2 Expression levels of putative genes with the function of glutamate signaling components in RNAseq analyses of maternal head (A) and embryo (B).** Gene expression levels are presented as the absolute values of Log_2_ (fold change) times –Log_10_ (*P* value). Genes differentially expressed between solitary and physical contact treatments are highlighted by red dots with the criteria of the absolute value of Log_2_ (fold change) > 0.58 and *P* value < 0.05. vGluT2 (LOC100168565) and mGluR3 (LOC107882629) were the only differentially expressed genes in the head and embryo, respectively. GS, glutamine synthetase; GltS, glutamate synthases; vGluT, vesicular glutamate transporters; NMDAR, NMDA receptor; GluRIB, glutamate receptor IB (AMPA receptors); KaiR1D, kainate-type ionotropic glutamate receptor subunit 1D; Ekar, eye-enriched kainate receptor; mGluR, metabotropic glutamate receptor.
