## Supplemental Figure S3 for "The maternal vGluT2 and embryonic mGluR3 signaling relay system controls offspring wing dimorphism in pea aphid"

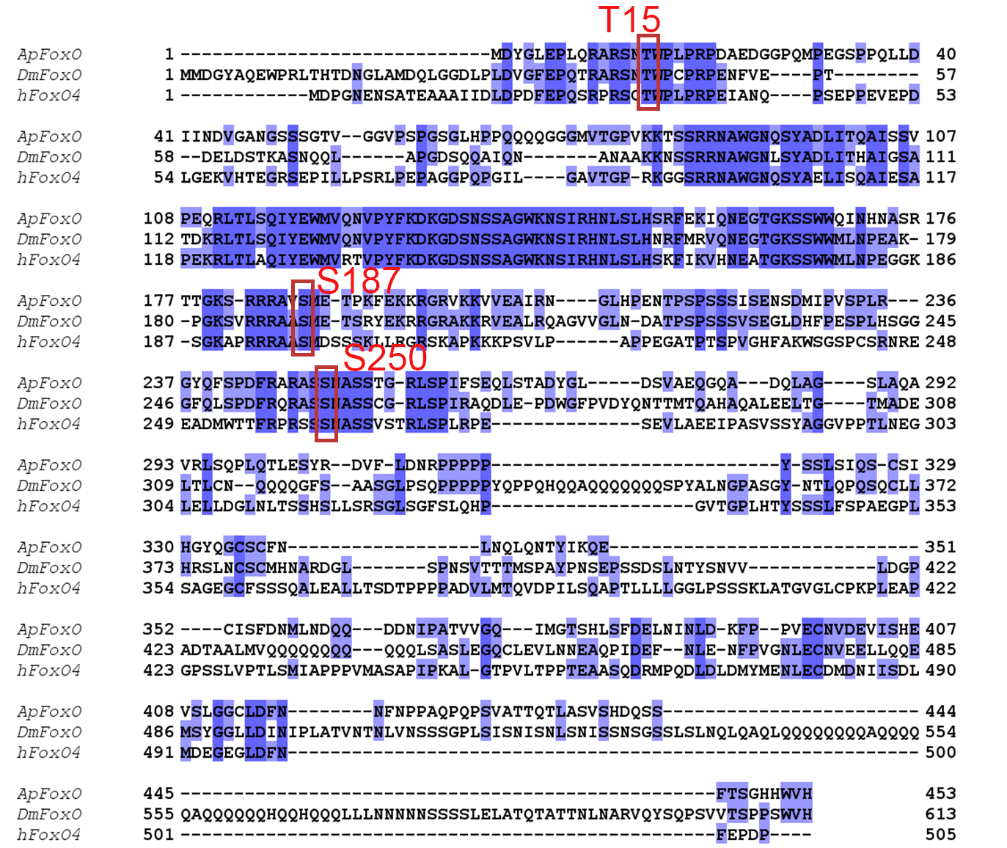

**Figure S3 Multiple amide acid sequence alignment indicating pea aphid FoxO potentially shares three conserved phosphorylation sites with *Drosophila* FoxO and Human FoxO4.** Three conserved phosphorylation sites, including T15, S187 and S250, were highlighted. *Ap*, *Acyrthosiphon pisum*; *Dm*, *Drosophila melanogaster*; *H*, *Homo sapiens*.
