## Supplemental Figure S4 for "The maternal vGluT2 and embryonic mGluR3 signaling relay system controls offspring wing dimorphism in pea aphid"

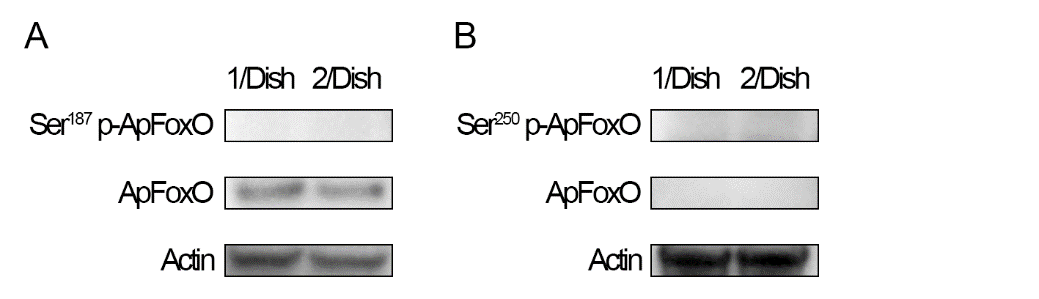

**Figure S4 Evaluation of phosphorylated *Ap*FoxO antibodies used in western blot targeting (A) phosphorylated S187-*Ap*FoxO, and (B) phosphorylated S250-*Ap*FoxO.**
