## Supplementary figures and images for "The maternal vGluT2 and embryonic mGluR3 signaling relay system controls offspring wing dimorphism in pea aphid"

### Supplemental Figure S5

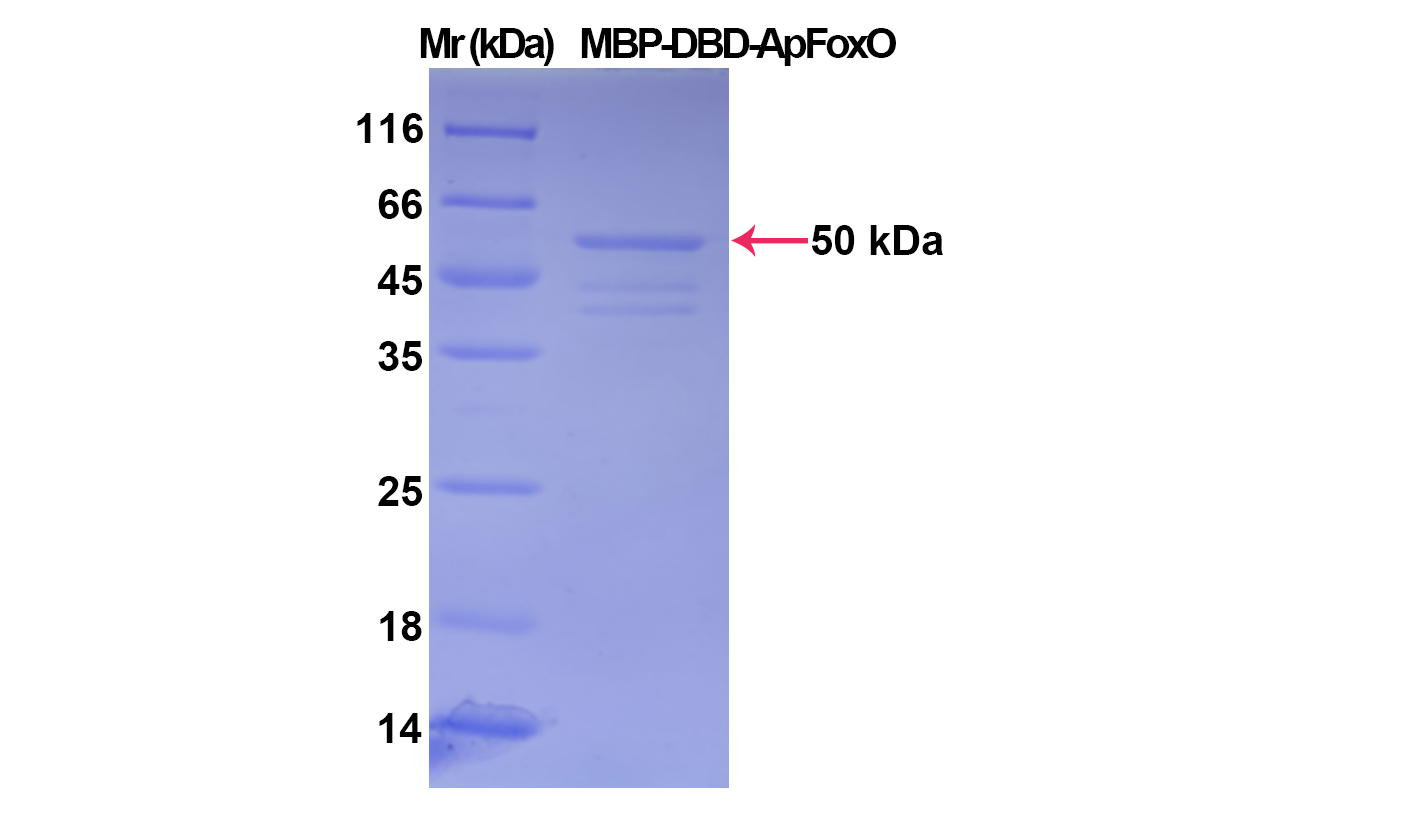

**Figure S5 SDS-PAGE of purified MBP--DBD_ApFoxO_ recombinant protein.**
