## Supplemental Figure S6 for "The maternal vGluT2 and embryonic mGluR3 signaling relay system controls offspring wing dimorphism in pea aphid"

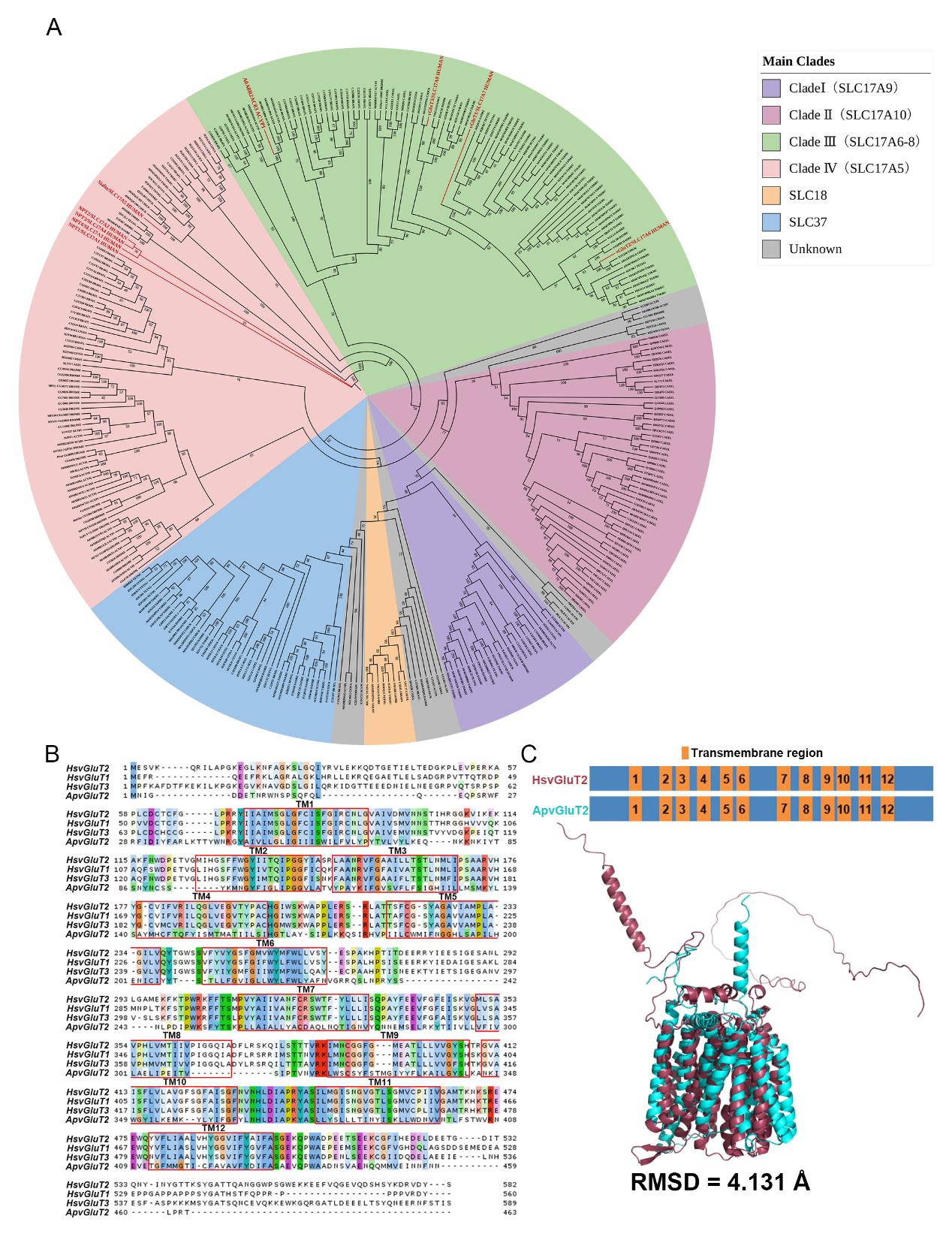

**Figure S6 ApvGluT2 is a structurally conserved protein with 12 transmembrane domains.** (A) The phylogenetic analysis of the vGluT2 family proteins indicated that ApvGluT2 belonged to the Clade III SLC17A9 subfamily. Bootstrap values expressed as percentage of 1000 replications were shown at branch points. Pea aphid vGluT2 (UniProt entry: A0A8R2AC83) and human homologs were highlighted by red characters. (B) Multiple sequence alignment and (C) pairwise structural alignment indicated that ApvGluT2 containing 12 transmembrane domains were similar to human homologs. Red rectangles marked 12 transmembrane domains. The structure of ApvGluT2 was predicted by AlphaFold2. The pairwise structural alignment was performed by PyMOL software. TM, transmembrane domain. ACYPI, *Acyrthosiphon pisum*; BRAFL, *Branchiostoma floridae*; CAEEL, *Caenorhabditis elegans*; CHICK, *Gallus gallus*; CIOSA, *Ciona savigyni*; DANRE, *Danio rerio*; DROME, *Drosophila melanogaster*; HUMAN, *Homo sapiens*; TAKRU, *Takifugu rubripes*; TETNG, *Tetraodon nigroviridis*. RMSD, root mean square deviation.
