## Supplemental Table S1 for "The maternal vGluT2 and embryonic mGluR3 signaling relay system controls offspring wing dimorphism in pea aphid"

**Table S1 Primers for RT-qPCR**

| **Gene name** | **Gene ID** | **Forward** | **Reverse** |
| --- | --- | --- | --- |
| RPL27 | ACYPI000085 | 5'–TCGTTACCCTCGGAAAGTC–3' | 5'–GTTGGCATAAGGTGGTTGT–3' |
| vGluT2 | LOC100168565 | 5'–AGGTTTAAGGATCAGCCCGC–3' | 5'–GCCCCAAGATCCCTGGAAAT–3' |
| mGluR3 | LOC107882629 | 5'–CCTCATATTGGACGACTGCGA–3' | 5'–ACGCCTTTCGAAGGTTCTGT–3' |
| GluCl-α | LOC100162577 | 5'–CCATGAACGTTTGGGATGGC–3' | 5'–ATTGGTAACGGCCAGTCTGA–3' |
| Octβ1R | LOC100166522 | 5'–AGTACGCTGTCGTGTCTTCC–3' | 5'–GTGCTTGTTCAGTAACGCGG–3' |
| GLRA1 | LOC100169196 | 5'–GGGTCCAACAAAGTTCACGC–3' | 5'–GGTCCACTGTTCGTCGTCAT–3' |
| HTR2A | LOC100575043 | 5'–GGTCAGACGAGACGGACATC–3' | 5'–GAATCGCCGTGACCCTTTTG–3' |
| FoxO | LOC100168097 | 5'–ACAGTTCGGCTGGATGGAAG–3' | 5'–GGCTCGCGTTGTGGTTTATC–3' |
| Dpp | LOC100163532 | 5'–GTCGGTTGGGACGATTGGAT–3' | 5'–TGGTGGAGTTGAGGTGTTCG–3' |
| Hh | LOC100165590 | 5'–GCGGGAGTGAAGAAGCAGAT–3' | 5'–CACGTTACGGTCCAGGAACA–3' |
| Vg | LOC100570322 | 5'–ACCAACAAGCGTCCACATCT–3' | 5'–CTGCTGTGGTGTGAGTCTGT–3' |
| Mats | LOC100160461 | 5'–CCACTCTTGGTTCGGGCAAT–3' | 5'–CCTCGGTGCAGAACTCTGTT–3' |
