## Supplemental Table S2 for "The maternal vGluT2 and embryonic mGluR3 signaling relay system controls offspring wing dimorphism in pea aphid"

**Table S2 Primers for dsRNA synthesis**

| Primer name | Gene ID | Forward | Reverse |
| --- | --- | --- | --- |
| GFP | - | 5'–CACAAGTTCAGCGTGTCCG–3' | 5'–GTTCACCTTGATGCCGTTC–3' |
| vGluT2 | LOC100168565 | 5'–AGTGCTCCTATCCTCCACGA–3' | 5'–AATATCGCGGGTTCAGGCTC–3' |
| mGluR3 | LOC107882629 | 5'–GTAGTCGAGGGCGATTTGCT–3' | 5'–ACAGATGTGACACTTGACGC–3' |
| FoxO | LOC100168097 | 5'–TTTCTCAGCCGCTACAGACG–3' | 5'–GTCATCCTGCTGGTCGTTCA–3' |
